# Programming Continuous, High Framerate, and Long-Duration Data Acquisitions on the Verasonics Ultrasound System

**DOI:** 10.64898/2026.11.25.769265

**Authors:** Matthew R. Lowerison, YiRang Shin, Yike Wang, BingZe Lin, Miguel Bernal, Bryan Cunitz, Jacques Battaglia, Jonathan Poree, Ronald E. Daigle, Olivier Couture, Jean Provost, Pengfei Song

## Abstract

The ultrasound community has seen a rise in popularity of highly demanding imaging acquisition sequences, such as those for super-resolution ultrasound localization microscopy and functional ultrasound imaging, which push the boundary of ultrasound system hardware limits. However, subtle errors in sequence programming can lead to corrupted data, inaccurate timing, and slow data throughput, which impact the performance of these modalities. In this article, we detail the fundamentals of the hardware and software sequencers for the Verasonics^®^ Vantage/NXT research ultrasound scanner, highlighting some common pitfalls in sequence design and ways to overcome them. We include example imaging sequences for different data acquisition goals, and detail recommended ways to test these sequences to ensure that they are meeting the expected performance.

## 2 Introduction

Over the last two decades, ultrafast and high data rate imaging has become a mainstream research tool in the ultrasound community [1]. Ultrafast imaging is the foundational technology for shear wave elastography, which requires a minimum imaging framerate of a few thousand Hertz (Hz) to adequately sample the typical frequencies that propagate in soft tissues [2–4]. These seminal works used planewave transmissions to cover the entire imaging area, and parallel processing with receive focusing to beamform images in software, which required a high transfer rate of raw radiofrequency (RF) channel data from the ultrasound hardware to the host workstation. Montaldo et al. [5] later formalized the process of coherently compounding multiple tilted planewaves to achieve transmit focusing, which yielded an image quality equivalent to conventional focused mode imaging at a ∼10 times faster imaging framerate, but increased the data transfer rate requirement. This high framerate tilted planewave strategy was leveraged to acquire large temporal ensembles to improve clutter filtering, resulting in ultra-sensitive Doppler performance [6], but at the expense of an even higher amount of data transfer.

During this time, fully programmable ultrasound research scanners with direct access to raw RF channel data, such as the Verasonics Vantage series, the Super-Sonic Imagine Aixplorer [6], the RASMUS system [7], and ULA-OP [8], to name a few, became widely available. This facilitated the development of highly demanding long-duration high framerate (LDHFR) imaging acquisition sequences. For example, functional ultrasound (fUS) [9] uses high framerate ultra-sensitive Doppler imaging, with the additional constraint of consistent imaging acquisition timing to accurately coincide with neurovascular stimulation patterns. fUS also requires long imaging durations, as the scale of neurovascular coupling can be several seconds, or longer, and sporadic/spontaneous studies require uninterrupted monitoring [10, 11]. Similarly, super-resolution ultrasound localization microscopy (ULM) [12, 13] demands high imaging framerate, large ensemble sizes, and long imaging durations to adequately scout microvascular flow [14–16]. These demands are compounded for functional ultrasound localization microscopy (fULM) [17], which combines the high data burden of ULM with the precise acquisition timing of fUS, or for dynamic ULM [18] which captures microvascular hemodynamics across several cardiac cycles. Translating these technologies from 2D to 3D, to achieve a larger field-of-view, less user dependence, and unbiased quantification of inherently 3D anatomy [19], further increase the demands on hardware and software.

It is common for LDHFR imaging sequences to require the acquisition, transfer, and storage of 10 to 1,000+ giga-bytes (GBs) of data. These acquisitions are often a “stress test” for imaging equipment as they frequently hit system hardware limits for both the host controller and the ultrasound research scanner, necessitating large local buffer sizes, high data transfer rates, high data writing speeds, and large data storage sizes. Next generation research ultrasound systems are also pursuing higher imaging frequencies and higher data sampling rates for small animal imaging [20], which further increases the data burden.

Critically, incorrect sequence programming and/or mismatched host controller hardware configurations can lead to corrupted RF channel data, avoidable bottlenecks that slow down data capture, and inaccurate acquisition timings (i.e., pulse repetition frequency (PRF), acquisition framerate, and fUS/fULM aggregate frame rate), which can substantially impact fUS and ULM/fULM performance. These errors can be subtle, often stemming from race conditions between the hardware and software sequencers: situations where both sequencers are attempting to access the same memory buffer simultaneously (e.g., the hardware sequencer is writing data while the host is attempting to read that data). This is particularly risky around direct memory access (DMA) events that transfer data from the hardware receive buffer into the host memory without continuous CPU involvement. Corrupted RF channel data can be difficult to diagnose and is often obscured after beamforming or in the presence of clutter.

For simplicity, we will be limiting our discussion to Verasonics sequence programming of coherently compounded planewave imaging acquisitions; however, the fundamentals described here are relevant to all ultrasound imaging systems and acquisition types. We are also focusing on saving RF channel data to disk as the main software-mediated process to navigate DMA events, as it is both broadly applicable and straightforward to demonstrate when there are errors. However, the principles described here also apply to other software processes that interact with an RF receive buffer, such as custom beam-forming implementations or digital filtering of RF data.

In this paper, we describe the fundamentals of the hardware and software sequencers of the Verasonics system and how they interplay for a typical LDHFR acquisition. We cover the basics of asynchronous and synchronous sequence programming, the benefits and risks of each, and provide working examples of LDHFR sequences. These sequences are available in a public GitHub repository [URL to be added when published] for dissemination to the ultrasound research community. Critically, these sequences provide utility functions that take the host hardware into consideration so that the user can make informed decisions about their acquisition parameters. Among these utility functions we include a sequence time-line visualization MATLAB app, an early pre-validation check that warns the user if there is a risk of RF corruption for a sequence under the given host configuration, and an RF timetag post-validation check to ensure that a user is meeting their stated acquisition goals. These utility functions are organized in Table 1 to help users identify what critical question in the sequence design and validation process the function is meant to address.

**Table 1:**
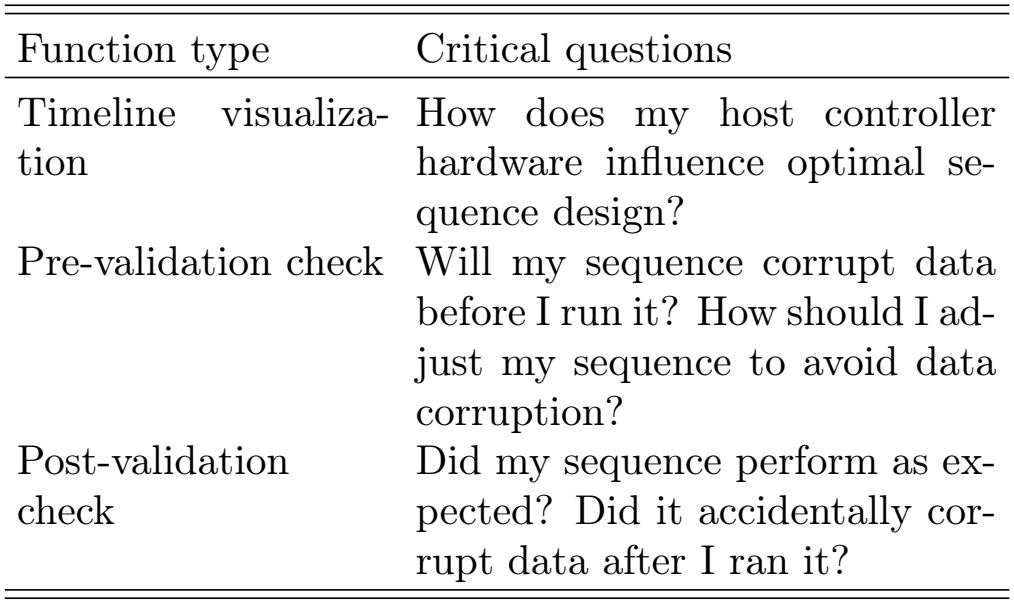
Sequence design utility functions.

| Function type | Critical questions |
|---|---|
| Timeline visualiza-tion | How does my host controller hardware influence optimal se-quence design? |
| Pre-validation check | Will my sequence corrupt data before I run it? How should I ad-just my sequence to avoid data corruption? |
| Post-validation check | Did my sequence perform as ex-pected? Did it accidentally cor-rupt data after I ran it? |

The rest of this article is organized as follows. In Section 3, we first describe the fundamental concepts and application of sequence design, including trade-offs in design space to meet imaging needs, and common pitfalls that may impact data quality. We also include strategies and methodologies for verifying that the sequence is performing as intended. We discuss data bottlenecks, make recommendations to maximize acquisition speed and through-put based on host hardware. In Section 4, we detail a case example applying one of these sequences to acquire a ULM dataset from a rat brain. In Section 5, we outline the results of this example imaging sequence, and the outcome of the sequence validation process. We also highlight the image reconstruction artifacts that can occur when RF corruption is present. In Section 6, we summarize our recommendations and observations from this case experiment and outline some limitations of the presented study.

## 3 Background

### 3.1 DMA, data race, and RF corruption

Before diving into specific details, it is necessary to introduce the terminology of an RF “super-frame,” which is a single memory buffer that contains the RF data from multiple transmit/receive events (e.g., the tilted planewaves for multiple imaging frames) into a contiguous data block. The general workflow of a LDHFR Verasonics sequence involves raw data capture of transmit/receive events into a local hardware memory buffer, a DMA transfer of this RF super-frame data into the host memory using a *transferToHost* (TTH) sequence control command, and finally a host software-mediated process to save this memory to disk for storage (**Fig. 1a**). The operator has freedom in selecting/designing the implementation of this saving process, which will be discussed in detail in Section 3.3, as it can have downstream effects for the optimal imaging sequence strategy. It is important to note that the hardware and software sequencers run independently of one another, and will both access the host memory for their respective events. The example diagram in **Fig. 1** has color-coded two RF super-frame acquisitions as blue and red to demonstrate this process. Careful coordination between hardware and software is essential to minimize the risk of data race conditions around DMA events, that is, situations in which a block of RF channel data is being simultaneously written to the host memory buffer by the hardware sequencer while being read from by the software sequencer (**Fig. 1b**). This can have deleterious and unpredictable effects on the quality of the RF channel data that is saved for later processing.

**Figure 1:**
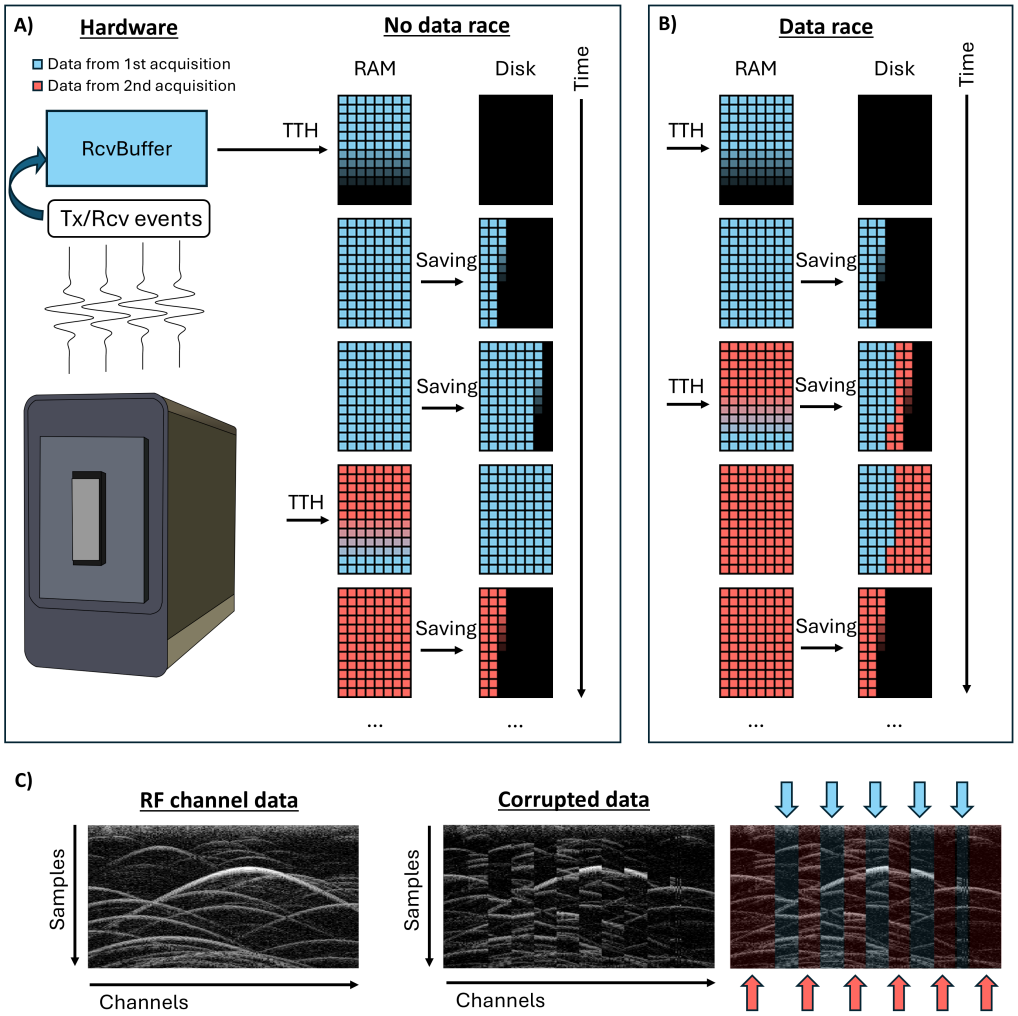
The hardware and software sequencers will independently access a receive buffer during an imaging acquisition. For this demonstration, we are color-coding the first RF super-frame acquisition in blue, and the second RF super-frame acquisition in red. **A)** In this example, there is no data race between the TTH and the software saving to disk, so the original RF channel data is intact. **B)** Example of a DMA data race, where the hardware sequencer begins transferring new data to the host memory buffer before the software sequencer has finished saving this data to disk. This results in a partial data overwrite which corrupts the RF channel data. **C)** Experimental data demonstrating a real-world example of data race associated-RF channel data corruption. The red and blue arrows highlight the mixing of channel data from two sets of transmit/receive events.

We will refer to DMA race conditions around RF channel data processing as “RF corruption.” An experimental example of RF corruption is shown in (**Fig. 1c**), which demonstrates data acquired from a beaker filled with degassed water mixed with microbubbles (MBs), with an intentional DMA race around data saving. The saved data contains a mix of two sets of transmit/receive events (denoted with red and blue arrows), which will negatively impact the overall quality of beamformed images. Due to the mixing of channel RF data from different time points there is also the risk of “phantom” MBs (i.e., MBs that are “retained” from the previous data acquisition), which complicates localization and tracking. A concrete example of a “phantom” MB is shown for an *in vivo* dataset in Section 5. This temporal ambiguity will also negatively impact functional imaging applications (e.g., fUS and fULM), leading to uncertainty in the correlation with stimulation patterns.

The most straightforward strategy for avoiding DMA race conditions is to intentionally synchronize the two sequencers (**Fig. 2**). This strategy sacrifices overall data throughput for a stronger guarantee of data integrity. Synchronizing the two sequencers can be accomplished implicitly by calling Verasonics image *recon* events to beamform receive buffer data, implicitly via a global *wait-ForProcessing* (WFP) flag (the preferred method in most circumstances), or explicitly using the *sync* command.

**Figure 2:**
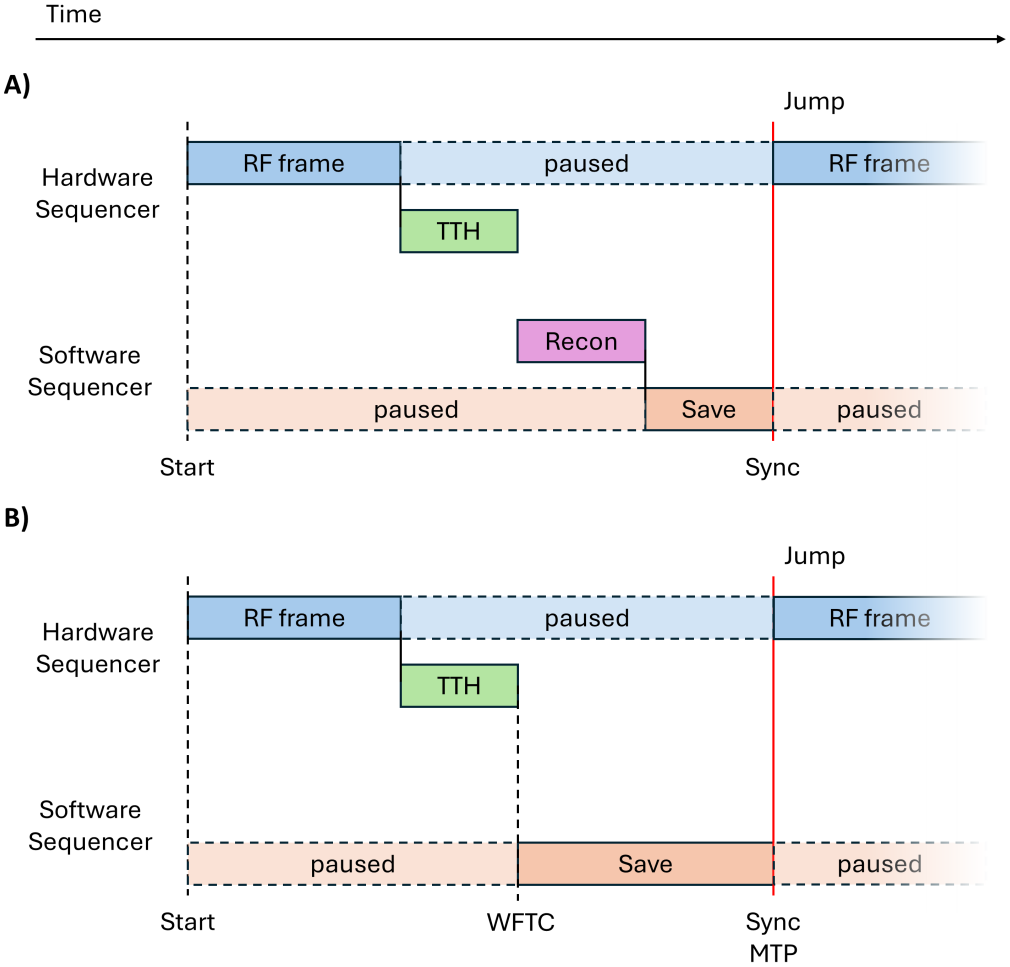
Example timelines of the hardware and software sequencers for synchronous data acquisitions. **A)** A call to Verasonics *recon* (i.e., beamform data) implicitly sets a *waitForTransferComplete* for the *transferToHost* event and then releases it with *markTransferProcessed* after reconstruction. **B)** An RF capture acquisition, which directly saves channel RF data for later processing. Critically, the *transferToHost* event needs to have a *waitFor-TransferComplete* set to prevent the software sequencer from jumping ahead too quickly. With this set up, the software sequencer will only begin saving RF data after the *transferToHost* is finished.

It should be noted that, for RF data acquisition sequences, merely including a *sync* command is insufficient to avoid a DMA race around TTH events: it is critical to either include a WFP flag to avoid a DMA race (discussed below), or a *waitForTransferComplete* (WFTC) command to temporarily pause the software sequencer until the entire receive buffer is transferred, as well as a *markTransferProcessed* (MTP) after saving to disk to ensure that every subsequent WFTC on these TTH events will be respected (e.g., for GUI controlled data RF acquisition sequence where every time a button is pressed it jumps to a set of events to save RF data and then returns to a live display loop).

The preferred synchronization strategy is the WFP flag, which provides the benefits of a strong guarantee of data integrity while also allowing for high data throughput and continuous acquisition under some circumstances. The WFP flag is a global toggle which ensures that the hardware sequencer does not advance more than one RF super-frame ahead of the software sequencer. As such, it is a good safety net in sequences which capitalize on alternating RF data buffer transfers (such as the concurrent design discussed in detail in Section 3.4) to ensure that there is no RF corruption occurring in saved data. It should be noted that WFP itself does not guarantee a continuous data acquisition; WFP will pause the hardware sequencer if the software sequencer is falling too far behind, but it will alleviate data corruption concerns for individual RF super-frames (**Fig. 3**). If the saving speed is faster than the RF data capture, then a WFP sequence will allow for a continuous (gap-less) data acquisition.

**Figure 3:**
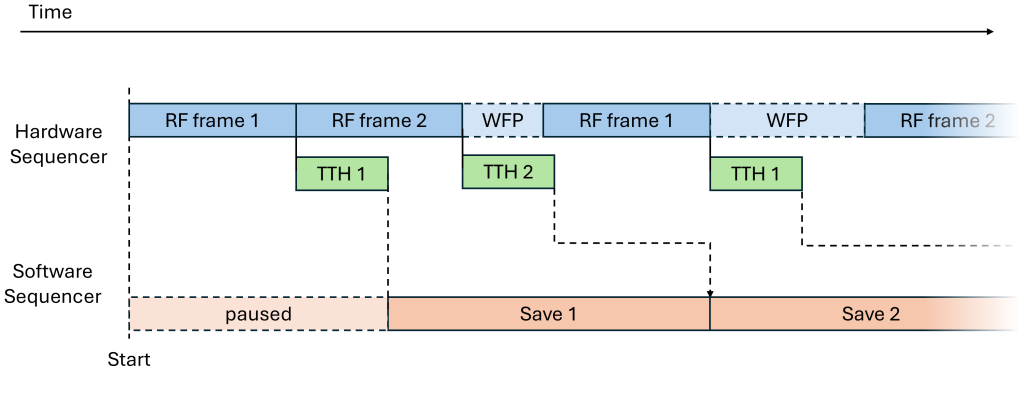
Example timeline of the hardware and software sequencers for synchronous data acquisition with a *waitForProcessing* flag and a long data saving time. In this specific example, the channel RF data saving process takes more time than the RF data capture, so the hardware sequencer slows down to accommodate the software sequencer, thus guaranteeing that no RF corruption occurs. In the situation where the saving process is faster than the data acquisition, this sequence would allow for a continuous (gap-less) data acquisition.

Although the strategy of explicit synchronization between the hardware and software sequencers is a good guard against RF data corruption, it has a few caveats. We are in some cases intentionally slowing down the data throughput, which exacerbates the long imaging acquisition times for ULM and fULM. Sequences with precise timing, such as fUS and fULM, may be less reliable if the software sequencer is dictating data capture, as there can be variability in the time taken to save or beamform data. It is preferred that fUS/fULM sequences instead rely on the hardware sequencer, as it has more consistent timing. To fully capitalize on the capability of the independent hardware and software sequencers we need to leverage asynchronous acquisition design. But before discussing strategies for designing around DMA transfers in asynchronous sequences, we need to outline a universal acquisition metric to fairly compare, and tune, the high variety of imaging sequences that are used in ultrasound research.

### 3.2 Data acquisition rate as a universal metric

Programmable ultrasound imaging systems provide a high degree of freedom for sequence design, the choice of transducer, imaging acquisition goals, etc. However, this complicates discussion of acquisition design as there are many factors that tend to be considered simultaneously. Even within the relatively narrow scope of coherent plane wave compounding a non-exhaustive list of parameters that a user may adjust includes: the number of samples per transmit/receive event (or equivalently the imaging depth relative to the transducer center frequency), the sampling rate, the number of channels, the number of compounding angles, ensemble size, and the final, intended imaging framerate.

Within the context of ensuring that there is no DMA data race we propose that the **data acquisition rate**, expressed in terms of GBs per second (GB/s), is a universal metric for sequence tuning (**Fig. 4a**). This allows us to consider sequence design within the context of two potential bottlenecks (**Fig. 4b**), one around the DMA transfer rate, and one around the saving speed of the host system. It also makes it straightforward to predict how adjustments to the sequence will impact the data acquisition rate, and therefore the risk of DMA race conditions. For example: if the number of compounding angles is doubled, and everything else is kept fixed, then the data acquisition rate doubles. If the imaging framerate is halved, then the data acquisition rate is halved. Understanding the relationship between the data acquisition rate and these two data throughput bottlenecks is critical for robust imaging sequence design. A sequence “timeline visualization” helper tool is provided as a MATLAB app in the GitHub repository in the “Benchmarking” subdirectory to facilitate this process.

**Figure 4:**
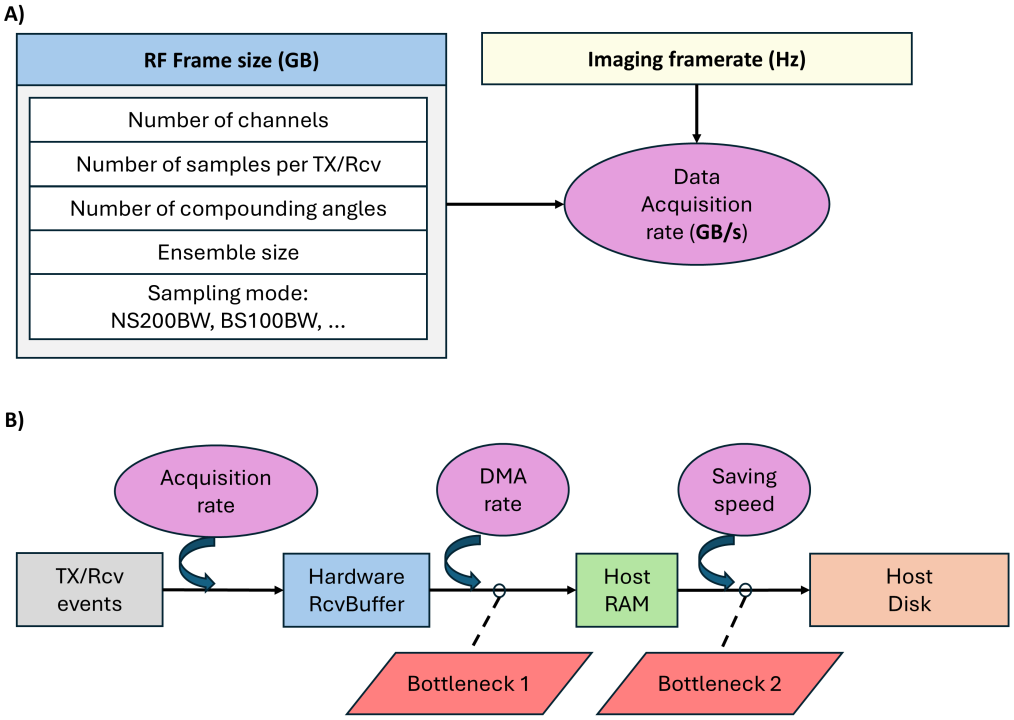
**A)** The data acquisition rate consolidates a wide range of user selected imaging parameters into a single metric. **B)** Outline of the data flow through for a Verasonics imaging sequence, highlighting two potential bottlenecks that need to be considered. The data acquisition rate sets the rate at which the hardware receive buffer is filled. A *transferToHost* call moves this data onto the host memory at the DMA transfer rate. The host configuration’s saving speed then determines the second bottleneck for safely saving this data onto disk.

### 3.3 Benchmarking bottlenecks

Consider the two main bottlenecks: the DMA transfer rate and the data saving rate. The DMA transfer rate is the absolute threshold for data acquisition rate that is achievable for an imaging sequence; under all circumstances it cannot be exceeded. Direct benchmarking of the DMA transfer rate is only available with the Verasonics *GPUDirect* toolkit, but it can be approximated using synchronized sequences with RF timetags (discussed in Section 3.5). There are also a few rules-of-thumb that can be applied to give a good rough estimate. The current generation of NXT uses acquisition boards with 8-lane PCIe Gen 3 connection, providing a bandwidth of roughly 7.1 GB/s, or 6.6 GiB/s. Some multi-connector univeral transducer adapters (UTAs), for example the 260-D UTA, split these lanes across the two connector ports, resulting in half the DMA bandwidth when using a 128-channel array. It should also be noted that the maximum DMA transfer rate is only achieved when the DMA buses are fully saturated, which occurs around an RF receive buffer size of greater than 0.1 GB (usually the case for LDHFR acquisitions). The DMA transfer rate is set by the NXT hardware, and thus there is little that a user can do to adjust it. However, there are edge cases where a misconfigured host controller can limit the DMA transfer rate: if the Verasonics adapter card is plugged into a host with a lower generation PCIe connector, or if that PCIe connector has fewer than 8 active lanes, then the DMA transfer rate will be slower than expected.

Benchmarking the saving rate is more straightforward, as it is defined by the user in software, but it is also more nuanced as it depends on the specific implementation, the host hardware and operating system, the file system, the type of hard drive/SSD/NVMe, how full the disk is, and so forth. These are discussed in more detail in Section 6. One important note is that most modern solid state data drives (SSD, NVMe, etc.) have a single-level cell (SLC) cache space that is used to temporarily store saved data at a higher speed before moving it to more permanent storage during drive idle time. During high data throughput this cache space can become exhausted, leading to a notable drop in speed to the sustained write rate.

In most situations, the user is defining a MATLAB external function in their Verasonics sequence to save the RF data (these can include MATLAB’s native *save* function, a call to *fwrite*, or a custom *mex* function), or they are using the Verasonics binary saving process command, and are often considering the trade-offs between write speeds, file size, compression rates, and compatibility. Another consideration is whether the saving process is blocking (i.e., the software sequencer waits until the save function returns, this is the case for most MATLAB implementations), or if it is leveraging multithreaded asynchronous file IO to avoid blocking (e.g., the Verasonics *fastSave* mex function). Asynchronous file IO is better able to take advantage of the fast SLC cache space of solid-state drives by distributing the save process across multiple drives, providing a larger amount of drive idle time for data consolidation. It should be noted that, for this technique to work, multiple solid-state drives are required for a single host, which increases the monetary expense of the system.

For benchmarking, it is suggested that the user wrap their saving implementation with MATLAB’s *timeit* function to provide a robust estimate of the actual time taken to save a hypothetical RF dataset of a known size. An example of this process is provided in the GitHub repository in the “Benchmarking” subdirectory. The Verasonics *fast-Save* mex function, which is available from the Verasonics Community Portal, provides an example benchmarking script for estimating the saving rate achievable using this external process.

### 3.4 Asynchronous acquisitions

With the data acquisition rate, DMA rate, and the saving rate on hand, we can now introduce the two general strategies for asynchronous acquisitions: one that capitalizes on a high data saving rate relative to the acquisition rate, and one that uses a large the host RAM allocation as a temporary buffer space for large continuous ensembles prior to a single block save. A decision tree for the selection of either of these two sequence types is shown in (**Fig. 5**), focusing on the option which gives the highest data throughput.

**Figure 5:**
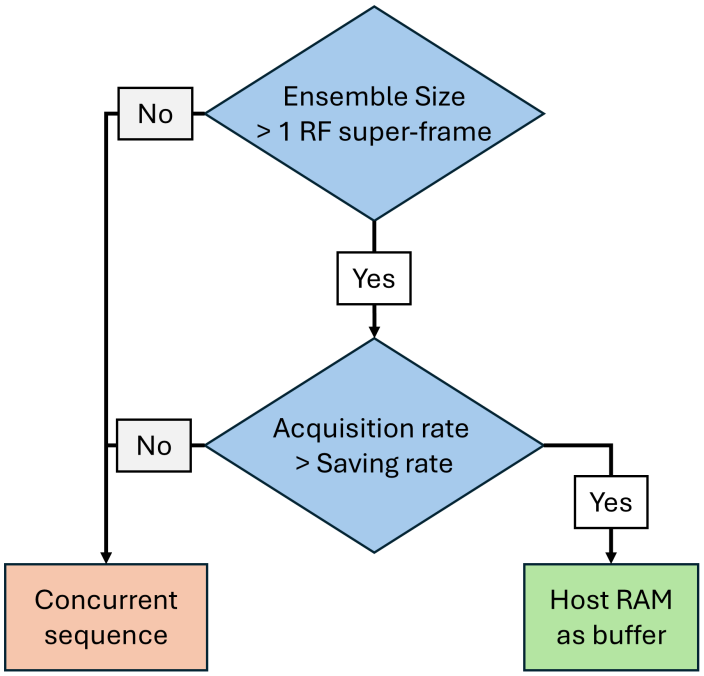
A decision tree for the selection of sequence strategy based on the bottleneck for a particular host configuration. If the data acquisition rate exceeds the rate that data can be saved, then the suggested strategy is to use the host memory/RAM as a temporary buffer space to allow large imaging ensembles, at the expense of a hardware pause between these ensembles. If the host can save data faster than the acquisition rate, then a concurrent imaging sequence is suggested.

The first consideration for sequence design is whether the total desired continuous ensemble size can fit within the 2 GB receive buffer limit for an RF super-frame. If this is the case, then a concurrent sequence is suggested, as it should be able to achieve the highest data throughput. This style of sequence is sometimes referred to as a “ping-pong” acquisition as the TTH transfer and *RF saving* buffers alternate to avoid DMA conflicts. An example of such a timeline is shown in (**Fig. 6**), where each saving process writes a single RF super-frame per function call. This is typical of fUS acquisitions, where discrete high framerate ensembles of a few hundred image frames are acquired at a set interval. These discrete ensembles are then clutter filtered and accumulated to generate a series of power Doppler images with an aggregate framerate determined by the gap interval. This is demonstrated on the timeline, where after each set of transmit/receive events into a single RF super-frame an additional *time-ToNextAcquisition* (TTNA) delay is added to the hardware sequencer, referred to here as a “long TTNA,” in order to make the timing of these relatively short ensembles consistent.

**Figure 6:**
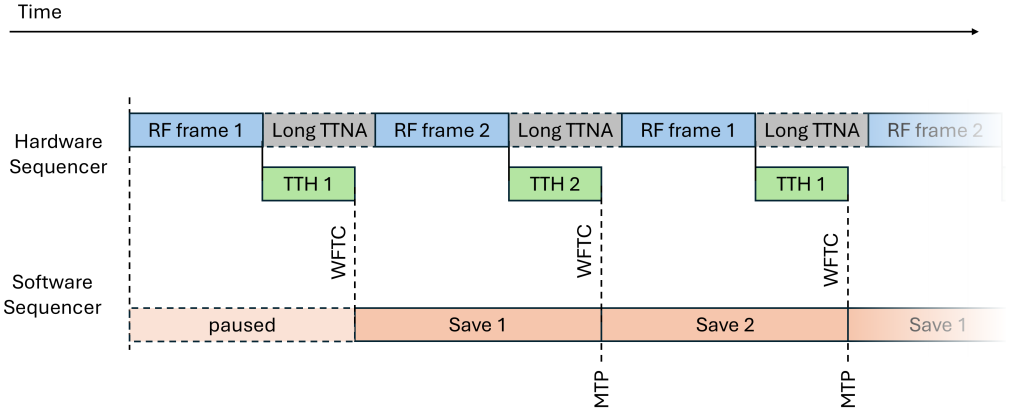
A concurrent-style asynchronous sequence. The saving process can overlap with the *transferToHost* event without risk of DMA race, provided that they are not referencing the same RcvBuffer data. The aggregate frame rate is set by the hardware sequencer with a long *time-ToNextAcquisition* at the end of each ensemble.

If the desired ensemble size is larger than what can fit into a single receive buffer frame, then the next consideration is whether the data acquisition rate exceeds the saving rate. If the saving rate is faster, then a truly continuous data acquisition (i.e., gap-less) is possible, and again a concurrent style of data acquisition is recommended. A timeline is shown in (**Fig. 7**). Continuous data acquisition allows for retrospective concatenation of arbitrarily long ensemble sizes, which can improve the performance of clutter filtering, especially for slowly flowing MBs in capillaries [21].

**Figure 7:**
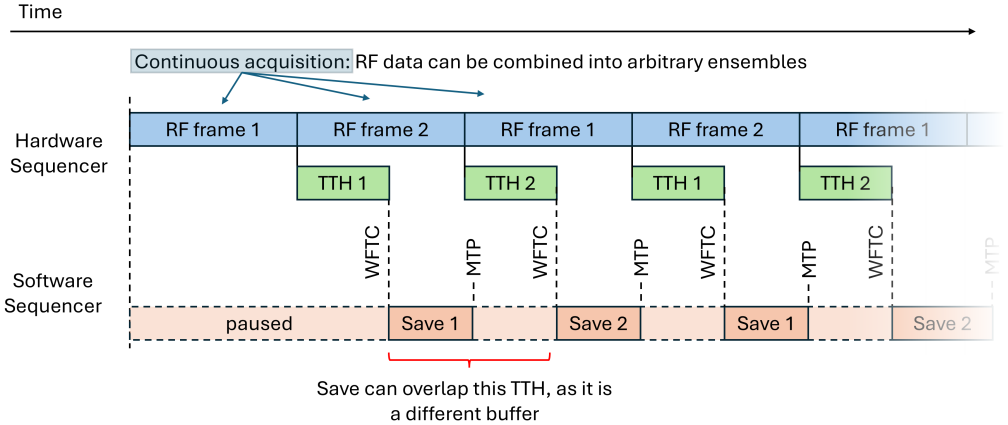
A continuous (gap-less) concurrent-style asynchronous sequence. As with the previous asynchronous example, the saving process can overlap with the *transferToHost* event under the same conditions. In this scenario, the saving rate is fast enough to allow the hardware sequencer to acquire data without gaps, allowing for arbitrarily long ensembles to be concatenated retrospectively.

The Verasonics *fastSave* mex function is designed to save a single RF super-frame per function call, leveraging asynchronous file IO and multiple save drives to maximize the SLC cache efficiency. The general sequence designed using the *fastSave* function is therefore a specialized version of a concurrent sequence, where the distributed write across multiple drives results in an effective saving rate that is faster than what a single drive could sustain. A timeline is shown in (**Fig. 8**). In this example, the software sequencer returns almost immediately, pausing only long enough to spawn a new thread for data saving that runs in the background. The data saving is distributed across multiple non-blocking drives, which can overlap, allowing for a higher effective saving rate than is possible under different drive configurations (e.g.: RAID0). In many scenarios there can be enough spare software sequencer time to do additional processing, such as beam-forming a few frames for a “live” update of imaging during an RF channel capture.

**Figure 8:**
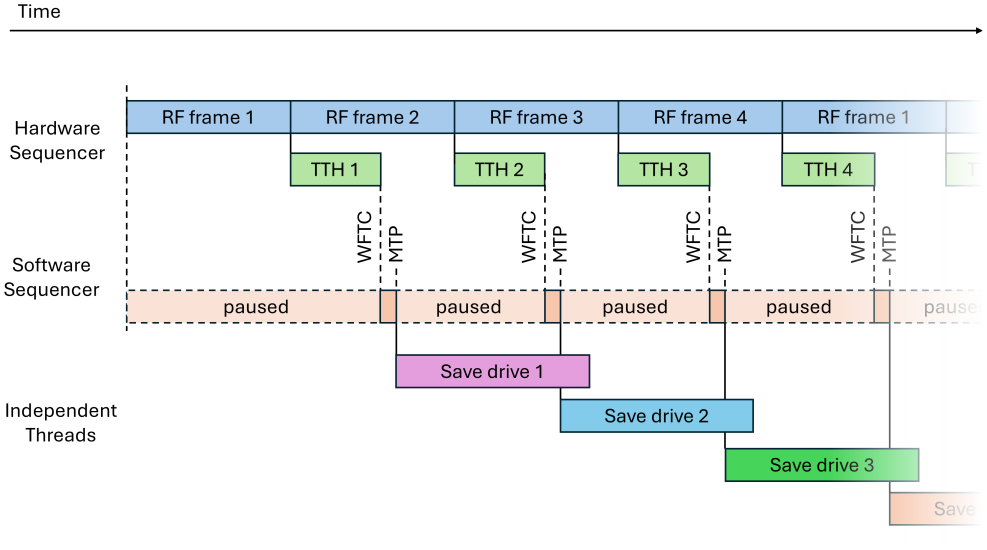
A continuous concurrent asynchronous sequence using Verasonics fastSave. The fastSave mex function distributes data writing across multiple data drives in a non-blocking manner on a separate independent thread, allowing the software sequencer to return almost immediately. This allows for a faster “effective” saving rate and frees up the software sequencer for other processes (e.g., beamforming a few image frames for live display).

The final situation involves a large ensemble size and a saving rate that is slow relative to the RF data acquisition rate. In this scenario, the host RAM memory can be used as a temporary buffer space to accumulate a large amount of channel RF data and then initiate data saving after a point where the risk of DMA race is minimized. A timeline is shown in (**Fig. 9**), where the saving process is writing multiple RF super-frames to disk. The “whole-block” data saving and TTH events can overlap to some degree (assuming that a subsequent call to a TTH event does not conflict with its respective portion of the saving block), but it should be noted that this style of data acquisition poses the highest risk of RF corruption: if at any point the software sequencer slows down then there can be a DMA race. If there is no need for precise timing between these large ensembles, then a multiple super-frame synchronous sequence should be preferred.

**Figure 9:**
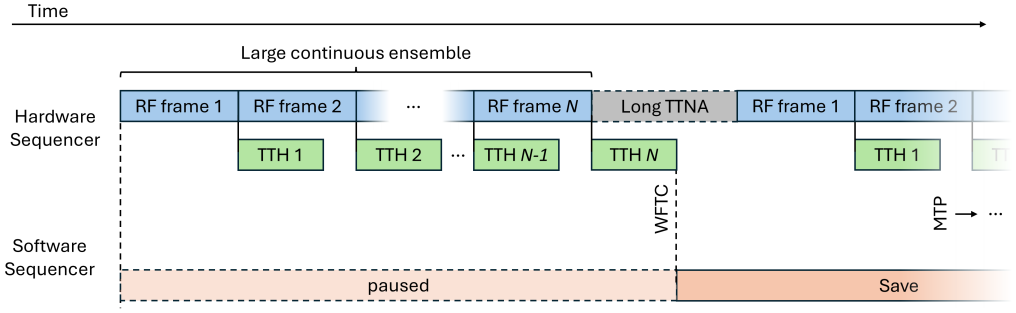
A RAM buffer asynchronous sequence. Multiple RF super-frame data acquisitions are acquired and transferred to the host memory, which serves as a temporary buffer space before starting the save process. This strategy is best when the saving rate is slower than the data acquisition rate as it allows for a longer continuous ensemble size.

### 3.5 Data Verification

Following RF data capture, there are a few steps that are suggested for ensuring that the imaging sequence was able to attain the requested data acquisition rate specification. The suggested approach is the use of RF time tags. The *Resource*.*VDAS*.*acqTimeTagEnable* command can be added to a sequence to write a 32-bit hardware timestamp into the first two samples of RF data in each system channel for each transmit/receive event. These timetags are set by the hardware sequencer but are only saved to disk by the software sequencer, providing a definitive answer to whether there was a DMA race occurring during data capture.

We have provided a helper function in the GitHub repository in the “Benchmarking” subdirectory for calculating and displaying this timetag data to determine if there was any RF corruption in a dataset. An example of this timetag analysis is shown later in Section 5 when we explore an experimental dataset acquired using one of the provided example sequences.

It should be noted that RF timetags only provide timing precision in increments of 25 *µ*sec, which is usually sufficient for between acquisition timing but may be inadequate for within acquisition timing. Users wishing to ensure that their sequence is meeting the specified TTNA for individual transmit/receive events are suggested to leverage the *trigger out* sequence control along with an oscilloscope to reach higher timing precision. However, there is an important caveat that the *trigger out* event is only characterizing the hardware sequencer, which does not guard against a DMA race when the data is being saved. Thus, it is suggested that *trigger out* hardware timing characterization is used as a supplement to RF timetag validation.

## 4 Methods

This section contains a case example of an RF capture experiment using the concurrent example sequence provided in the repository for ULM processing. The host controller was benchmarked using the provided repository tools to design a stable continuous (gap-less) acquisition.

### 4.1 Host controller

The host controller was a custom HP^®^ Z6 G5 A tower workstation running Ubuntu 22.04, with a 24-core AMD Ryzen^−^ Threadripper^−^ PRO 7965WX processor, an NVIDIA A6000 GPU, and 256 GB DDR5 onboard memory. For data storage it had a 4TB M.2 drive (Crucial^®^ T705) formatted with the F2FS filesystem, and a Dell ultra-speed quad NVMe bridge with four 2TB NVMe drives (Samsung 990 pro) formatted with the ext4 filesystem. Save rate benchmarking of these drives yielded an approximate MATLAB fwrite speed of 1.6 GB/s for the 2TB NVMe drives, and 3.2 GB/s for the 4TB M.2 drive.

The *fastSave* benchmarking script for the 2TB NVMe drives yielded an SLC cache write rate of 6.1 GB/s until the cache filled (around 210 GB written), at which point it dropped to a 1.5 GB/s sustained write rate.

### 4.2 Animals

All animal procedures were approved by the Institutional Animal Care and Use Committee at Duke University (IACUC Protocol number A028-25-04). All presented experiments were conducted in accordance with the IACUC guidelines. Every attempt was made to minimize the number of animals used and to reduce suffering at all stages of the study.

A 25-week-old Sprague-Dawley rat was used in this study to demonstrate the impact of RF data corruption on ULM reconstruction fidelity. The animal was anesthetized using 3% isoflurane mixed with medical oxygen for induction and kept at 1.5% isoflurane for surgery and imaging. Once under anesthesia, the animal was transferred to a stereotaxic imaging stage (RWD) and was secured via ear bars. A craniotomy was performed using a rotary Dremel^®^ tool to expose the brain, and a catheter was inserted into the tail vein for infusion of Lumason^®^ MB contrast agent (Bracco, Milan, Italy). This MB solution was infused at a rate of 50 *µ*L/min using a programmable syringe pump (NE-300, New Era Pump Systems Inc., Farmingdale, NY, USA).

### 4.3 Ultrasound Data Acquisition

All ultrasound data collection was performed using a Vantage^®^ NXT 256-channel research ultrasound system (Verasonics Inc., Kirkland, WA, USA). An L22-14vX (Verasonics Inc., Kirkland, WA, USA) linear array transducer was secured to the stereotaxic frame via a custom 3D-printed transducer holder and positioned for a coronal anatomical imaging plane.

Imaging was performed using the concurrent example sequence in the repository as a reference (**Fig. 8**), with the RF time tag option enabled. We modified the sequence to have 8-angle plane-wave compounding (2-degree increment). Other imaging parameters were set to a transmit center frequency of 15.625 MHz, a 1000 Hz post-compounding frame rate, a transmit voltage of 6 volts, and “BS100BW” sampling at a 15.625 MHz demodulation frequency. Each saved RF acquisition represented an ensemble size that could be beamformed into 500 IQ image frames, with 896 RF samples per transmit/receive event. This yielded an acquisition data rate of 1.84 GB/s, which is well within the benchmarked saving rate of the host controller used in this study, providing a continuous data acquisition. The total imaging duration was 180 seconds, and the final data size for this experiment was 330 GB. This channel RF data was beamformed offline using a custom CUDA-based delay-and-sum GPU beamformer.

### 4.4 ULM processing

All ULM processing was performed in MATLAB (The MathWorks, Natick, MA; version R2022b). Clutter filtering was performed using singular-value decomposition [22] with a low-order tissue threshold of 20 to extract the MB signal. A MB separation filter [23] was used to split the MB data into upward and downward flow to be processed separately. MB localization was performed using normalized cross-correlation with a 2D Gaussian template and a threshold of 0.6 [24]. MB tracking was done with a custom minimum-distance pairing algorithm, and MB trajectories were plotted onto a reconstruction grid space with a pixel size of 2 *µ*m. These trajectories were scaled based on the MB backscatter intensity for visualization [25, 26].

The estimated ULM resolution was calculated using Fourier ring correlation (FRC) [27], following the protocol detailed in [28], where each MB track is randomly assigned to one of two ULM sub-accumulations to produce independent reconstructions of the ULM dataset. The resolution was estimated by finding the crossing point of the FRC curve with two thresholds that are commonly used for ULM characterization: the 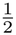-bit threshold and the 2-*σ* criterion [29].

### 4.5 Synthetic RF corruption

To investigate the impact of a hypothetical DMA race on ULM performance, we conducted a synthetic channel RF corruption experiment on the rat brain dataset from the previous section. The synthetic RF corruption was performed as follows: for every RF super-frame we retained a total of 32 randomly selected RF channels, in blocks of 8 contiguous channels, from the previous un-corrupted data acquisition. This was done to approximate the experimental RF corruption seen in **Fig. 1c**. The first RF dataset was left unchanged, and this synthetic corruption process was done independently for each subsequent dataset. This represents a corruption of approximately 25 % of the RF channel data, with a maximum temporal shift of 500 msec between datasets. The beamforming and ULM processing were kept the same between the original and corrupted RF datasets.

## 5 Results

### 5.1 Rat brain ULM acquisition validation

The proposed concurrent imaging acquisition sequence had a data acquisition rate which was below the bench-marked saving rate for the host controller, implying that continuous acquisition was possible. A visualization of this sequence, generated using the provided MATLAB app, is demonstrated in (**Fig. 10a**) as a continuous (gapless) concurrent sequence. It should be noted that this only includes a short section of the total proposed imaging duration (only around 8 seconds of the total 180-second imaging time), which was done for the purposes of visualization clarity in the figure.

**Figure 10:**
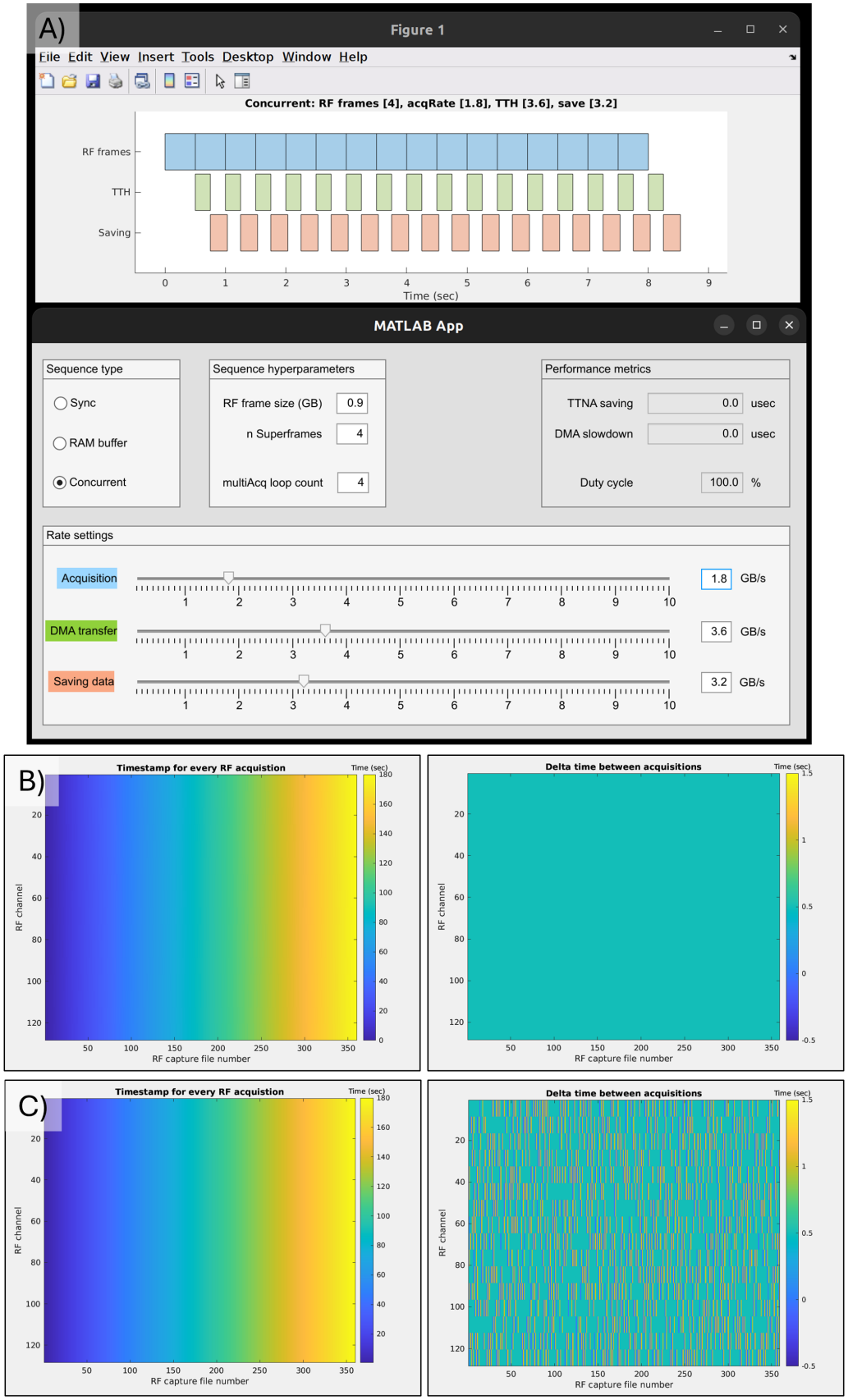
Output of some of the provided benchmarking tools for the example rat ULM imaging sequence. **A)** The MATLAB app-based acquisition timeline visualization tool predicted that the proposed sequence parameters would allow for continuous acquisition of channel RF data. **B)** RF timetag-based data validation. The time between every RF super-frame was consistently 0.5 seconds for all channel data for every acquisition. This indicates that there was no RF corruption in the example dataset. **C)** Example RF timetag output for the synthetically corrupted dataset, where the time between each channel varied from 0 to 1 second. This indicates a mixing of temporal information between RF super-frames.

After the imaging experiment, we conducted a post-validation of the RF timetags using the provided benchmarking tools. The output of this benchmarking is demonstrated in (**Fig. 10b**), which plots the delta time between all 360 RF super-frames that were acquired in this experiment. This indicates that there was no RF corruption in this dataset, and that it represents a true continuous dataset with a total duration of 180 seconds. For comparison, we also included the timetag output for the synthetic RF corruption data set (**Fig. 10c**), to highlight what a hypothetical sequence with a DMA race would look like with these benchmarking tools.

### 5.2 Impact of corruption on diffraction limited images

As mentioned previously, race conditions between the hardware and software sequencers around DMA events can be difficult to diagnose as the impact of RF corruption can be obscured by beamforming or in the presence of tissue clutter. This is demonstrated in (**Fig. 11a** and **b**), where the RF channel data and B-mode images of the datasets with and without RF corruption are perceptually nearly identical. Following SVD filtering (**Fig. 11c**) it is also difficult to determine if there was an issue. We can observe that the MB point-spread functions (PSFs) have been broadened in comparison to the no data race condition, particularly in the lateral direction, which will impact localization accuracy. This broadening of the PSF may also impact the localization efficiency, as nearby MBs which were spatially isolated may now overlap. This is demonstrated with the blue arrows in (**Fig. 11d**). We also have evidence of a “phantom” MB phenomenon, shown with the red arrow, which occurs when there is sufficient “stale” RF channel data from the previous acquisition polluting the current RF super-frame to reconstruct an MB-like imaging feature.

**Figure 11:**
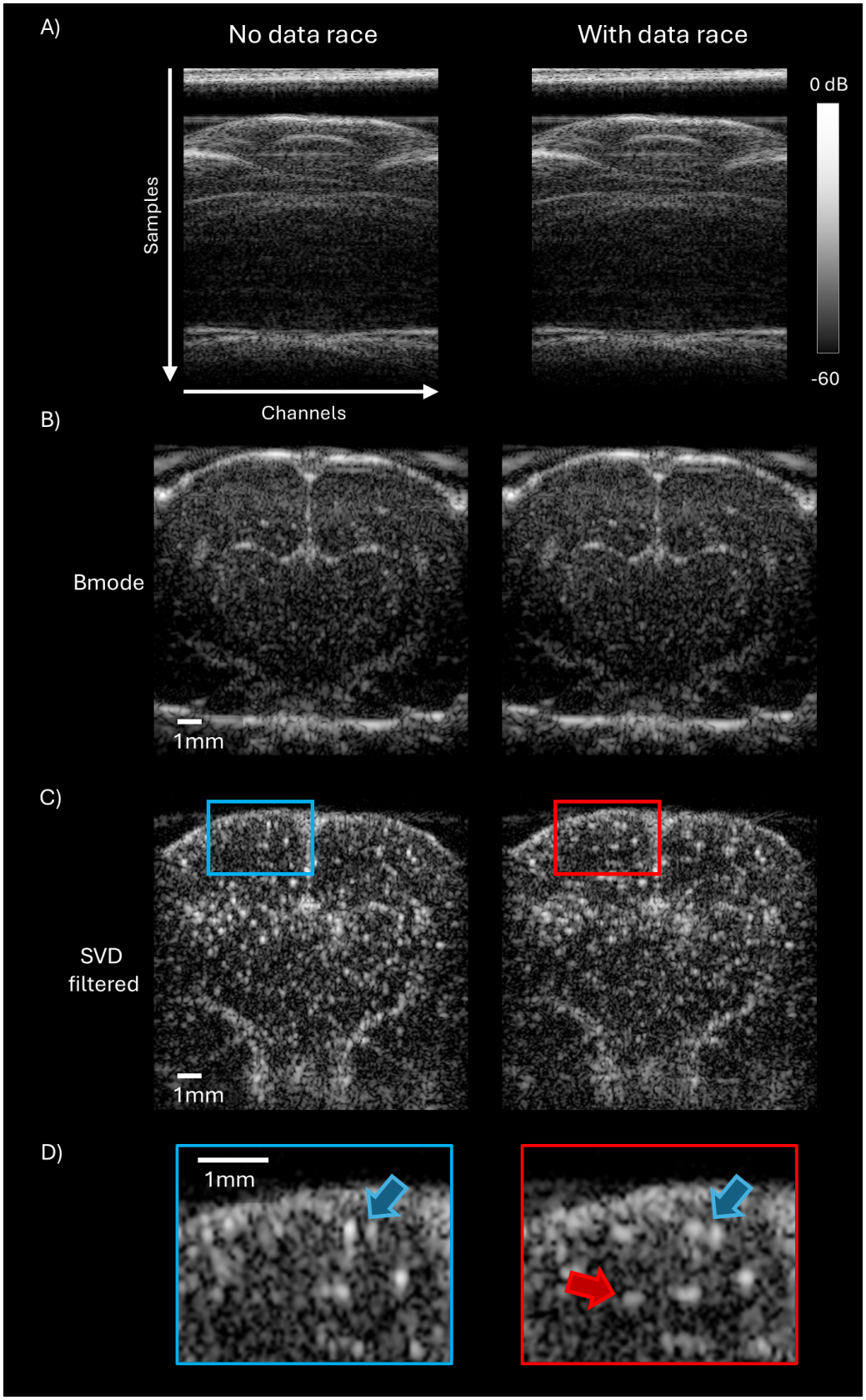
Comparison of the experimental rat brain ULM dataset under no data race and with data race conditions. **A)** Without timetag information, it can be difficult to discern if RF corruption has occurred in the presence of tissue clutter. **B)** Beamforming obscures RF corruption, with nearly identical B-mode images between the two datasets. **C)** SVD filtering reveals a subtle shift in the MB appearance, with wider lateral expanse of the apparent PSF. **D)** A zoomed in view of the cortical region demonstrates a higher risk of MB overlap (blue arrow) and a “phantom” MB in the corrupted dataset (red arrow).

### 5.3 Impact of corruption on ULM

A ULM reconstruction from the rat brain concurrent script example experiment, is shown in (**Fig. 12a**). We note that there is generally good microvascular reconstruction performance throughout the cortical, hippocampal, thalamic, and other deep brain regions. We selected a few regions-of-interest from these anatomical structures to highlight the impact that the data race has on ULM reconstruction quality. These are shown in the insets of (**Fig. 12b**), where we can note that the synthetic data race RF corruption led to blurring of the arterioles/venules in the ULM image, and many of the finer microvascular features are obscured.

**Figure 12:**
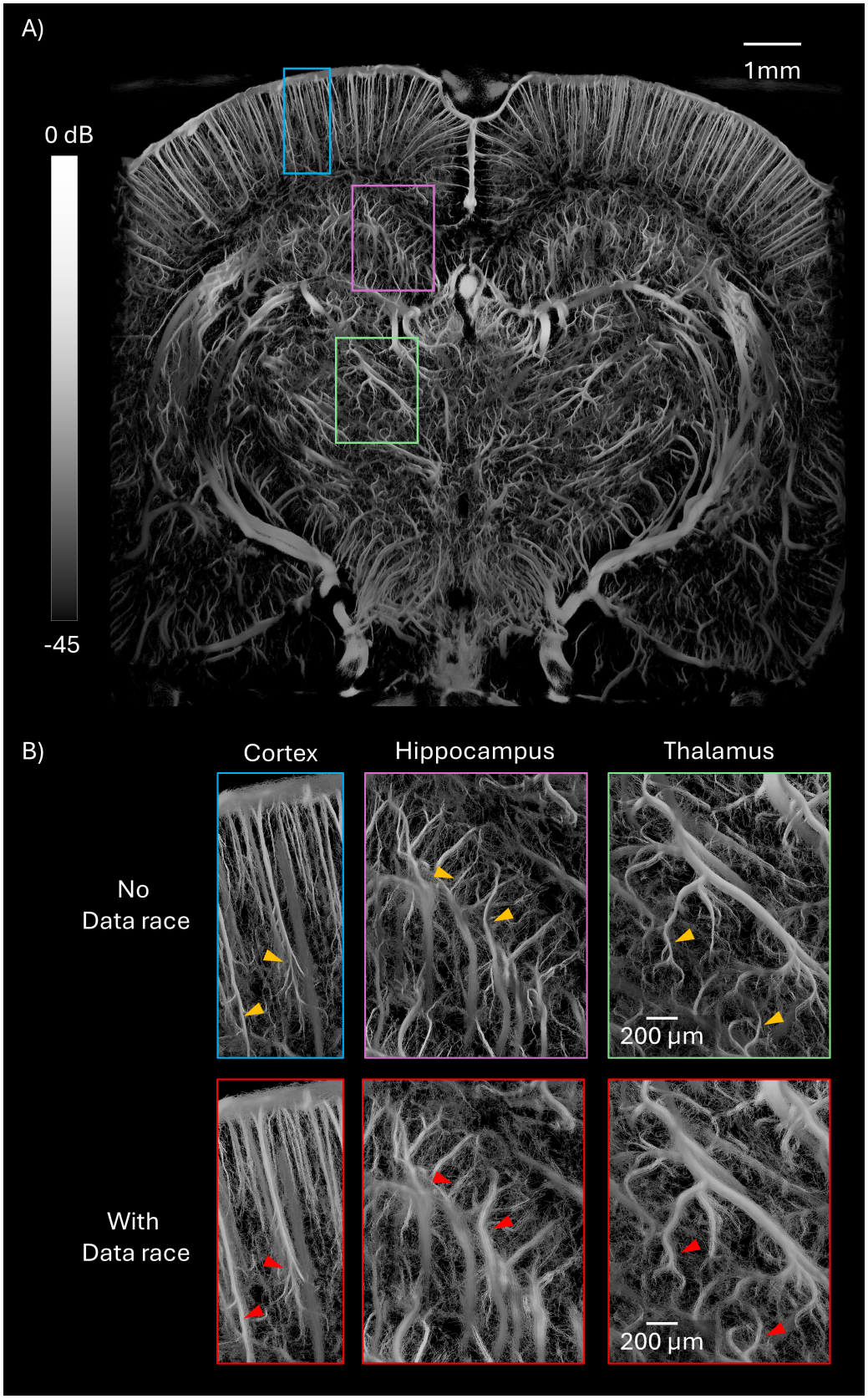
**A)** ULM reconstruction of the rat brain dataset acquired using a concurrent continuous acquisition with an L22-14vX. **B)** Zoomed-in insets of different anatomical regions highlighting the blurring effect of RF data corruption on ULM reconstruction performance, with arrows indicating a few case examples were microvascular detail is lost due to this effect.

This difference in ULM performance was quantified using FRC, as demonstrated in (**Fig. 13**). The base-line ULM reconstruction had an estimated resolution of 9.2 *µ*m according to the 2-*σ* criterion, which is approximately 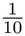 of the wavelength of the 15.625 MHz transmit center frequency (98 *µ*m for an assumed sound speed of 1540 m/s). The stricter 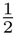 -bit threshold yielded an estimate of 16.2 *µ*m. The impact of synthetic RF corruption was profound, with a 2-*σ* resolution estimate of 16.5 *µ*m, and a 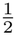 -bit estimate of 24.0 *µ*m. This could still technically be considered a super-resolved result; however, the impact on the quality and performance of ULM should not be understated.

**Figure 13:**
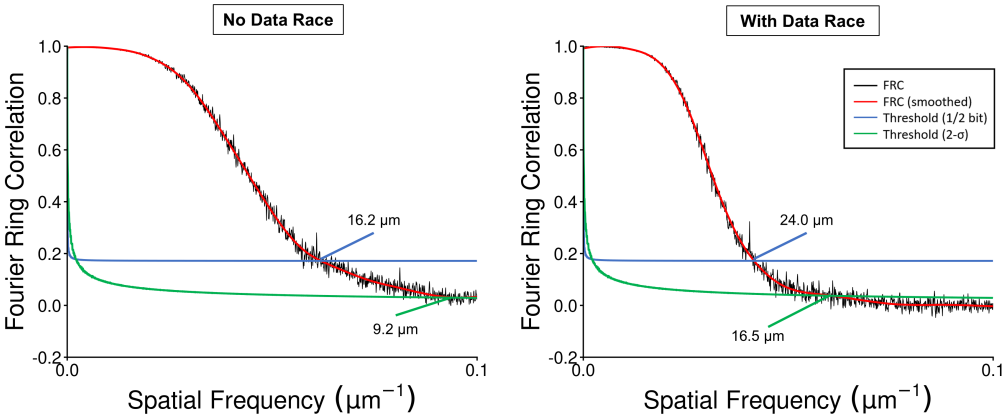
FRC analysis to estimate ULM resolution. The baseline example dataset yielded a resolution estimate of 9.2 *µ*m under the 2-*σ* criterion, which is below one-tenth of a wavelength. The same dataset under a synthetic data race condition substantially impacted ULM performance.

## 6 Discussion

Ultrasound imaging is a rapidly developing and dynamic field of research. The ultrasound community has embraced highly demanding imaging acquisition sequences, and novel ultrasound imaging development and applications are widespread. This community has been empowered by access to programmable ultrasound systems, such as the Verasonics NXT, Ultrasonix, or ULA-OP, which provide powerful and flexible platforms to conduct research. However, these systems also present a high degree of complexity and a substantial learning curve, which makes *de novo* imaging sequence design difficult and rife with potential errors that can impact the quality of re-search. These risks are inherent to the design and flexibility of these systems and thus should be considered carefully for the majority of ultrasound imaging research platforms.

One substantial concern that we have highlighted in this study is the risk of channel RF corruption stemming from race conditions around DMA data transfer events between the ultrasound hardware and the host controller. This form of RF corruption impacts LDHFR imaging performance (**Figs. 12, 13**) but paradoxically may be difficult to detect without careful examination and postvalidation strategies (**Fig. 11**). Furthermore, the case example of this RF corruption phenomenon presented in **Fig. 1c** is only obvious in the channel data as this experiment was for a simple imaging target: low concentration MBs in water. The addition of tissue clutter, noise, motion artifacts, and so forth for an *in vivo* imaging experiment can also obscure this RF corruption effect, as is demonstrated in (**Fig. 11a**).

To address these issues, we have proposed a universal metric for sequence tuning to avoid RF corruption, provided example imaging sequences with built-in prevalidation steps to warn the user if there is the risk of poor data quality, and outlined a post-validation workflow for ensuring that the RF data meets the expected data acquisition rate target. We have also highlighted the specific sequence control commands of the Verasonics platform for navigating around DMA data transfer. These include WFTC/MTP pairs as a general guard around RF TTH events to ensure that the software sequencer does not access RF data before the hardware has finished the transfer, and WFP as a safety net in concurrent sequences to avoid the hardware outpacing the software sequencer. These data integrity checks are critical for ultrasound imaging sequences generally, and become even more relevant for long duration imaging sequences which may span tens of minutes [10, 11, 17, 30–32].

An essential step of sequence design is the consideration of the host system, especially with respect to the data saving rate and the available system memory. To this end, we have provided benchmarking tools for estimating the saving speed bottleneck and summarized a simple decision tree (**Fig. 5**) to help users select which imaging sequence would be best suited for their specific equipment set-up. The provided example sequences also report a diagnostic output to the command window which outlines the RF buffer size, data acquisition rate, number of RF frames that can fit into the available host memory, and any additional TTNA which may be required for saving RF data after the acquisition of an RF super-frame ensemble. This additional TTNA serves as an indicator as to whether or not the current sequence parameters would allow for a continuous data acquisition.

This is meant to be a pragmatic solution for optimizing a sequence based on the current host controller, which is critical given the increasing costs of data storage and memory. Understanding these bottlenecks can also be used to better identify what upgrades to the host controller would provide the best return on investment for a specific research goal. It should also be noted that there are many factors which influence the data writing rate even with fixed hardware. In the host characterization portion of the methods, we demonstrated a case example where formatting an NVMe drive to the F2FS filesystem, which is better optimized for solid-state drives than ext4, achieved roughly double the write speed for MAT-LAB fwrite. The implementation of the saving function has a substantial impact on the data saving speed, where asynchronous file IO can better leverage SLC cache space. Data compression during RF saving can save disk space but will add computational overhead. Solid state drives also slow down the more of the storage space is occupied, so emptying a drive before attempting a long data acquisition is critical for ensuring reliable performance. These factors should be considered and optimized on a per-host basis.

This study has a few limitations that should be addressed. The synthetic corruption comparison presented in **Fig. 12** is an idealized case of a DMA race condition, with a set pre-determined number of “corrupted” RF channels that were delayed by exactly one RF super-frame. More realistically, a DMA race is stochastic and depends on the specifics of the sequence design and the current state of the host controller. The severity of the race condition would likely change over the course of a long imaging experiment, as the saving speed will slow down as the SLC cache fills, and any additional processing calls (e.g., from the operating system or other software running simultaneously) may change the timing of the software sequencer. We opted for a synthetic corruption process to mitigate some of this variance, and to provide a clean baseline result for a direct comparison of performance degradation due to a hypothetical DMA race.

The division of asynchronous data acquisition schema into either save rate mediated or host RAM mediated is a bit of an oversimplification. In reality, all LDHFR sequences use at least some host RAM as a temporary buffer space, and all LDHFR sequences must save data to disk. Hybrid sequences, for example a RAM buffer style acquisition which uses Verasonic’s *fastSave* mex function to save discrete ensembles exceeding the RF super-frame page size (e.g., 4 RF super-frames acquired continuously, but with a gap between each acquisition of 4 RF super-frames) are certainly possible. These were omitted due to the large scope and complexity required to adequately cover every permutation of hybrid sequence design.

Only one host configuration was thoroughly bench-marked during this study, and the system was more powerful than the default host controller supplied by Verasonics. We had the luxury of being able to clear all data from our saving drives before each experiment. This system was also running using a Linux-based operating system, which may have different performance metrics for asynchronous file IO compared with a Windows-based host. The principles should also apply to other host controllers and operating systems, but these were not rigorously tested during the development of the supplied sequences. Saving rate benchmarking was only performed on a limited subset of potential user implementations. Other considerations, such as data compression, different file formats, and alternative binary write implementations, should be explored more rigorously.

We also focused on RF timetag analysis as the main form of data validation for detecting DMA race conditions and therefore RF data corruption. As mentioned previously, RF timetags provide timing precision in increments of 25 *µ*sec and therefore may be too coarse for some timing characterization or they may mask a very slight DMA race.

## 7 Conclusion

In this study, we presented a framework for rigorously designing and validating continuous, high framerate, and long-duration data acquisition sequences on the Verasonics NXT ultrasound system. A set of example sequences were provided, along with benchmarking and testing functions, for dissemination to the ultrasound research community. Critically, we highlight a common pitfall associated with asynchronous data capture which can impact ultrasound imaging performance and provide concrete implementations that consider the host hardware to ensure that these errors are not encountered.

